# The uptake of tau amyloid fibrils is facilitated by the cellular prion protein and hampers prion propagation in cultured cells

**DOI:** 10.1101/2020.01.19.911644

**Authors:** Elena De Cecco, Luigi Celauro, Silvia Vanni, Micaela Grandolfo, Adriano Aguzzi, Giuseppe Legname

**Affiliations:** Laboratory of Prion Biology, Department of Neuroscience, Scuola Internazionale Superiore di Studi Avanzati (SISSA), Trieste, Italy; Institute of Neuropathology, University Hospital of Zürich Schmelzbergstrasse 12, CH-8091 Zürich, Switzerland; ELETTRA Sincrotrone Trieste S.C.p.A, Basovizza, Trieste, Italy

**Keywords:** prion protein, tau, prion propagation, internalization, scrapie

## Abstract

Tauopathies are prevalent, invariably fatal brain diseases for which no cure is available. Tauopathies progressively affect the brain through cell-to-cell transfer of tau protein amyloids, yet the spreading mechanisms are unknown. Here we show that the cellular prion protein (PrP^C^) facilitates the uptake of tau aggregates by cultured cells, possibly by acting as an endocytic receptor. In mouse neuroblastoma cells, we found that tau amyloids bind to PrP^C^; internalization of tau fibrils was reduced in isogenic cells devoid of the gene encoding PrP^C^. Antibodies against N-proximal epitopes of PrP^C^ impaired the binding of tau amyloids and decreased their uptake. Surprisingly, exposure of chronically prion-infected cells to tau amyloids reduced the accumulation of aggregated prion protein; this effect lasted for more than 72 hours after amyloid removal. These results point to bidirectional interactions between the two proteins: whilst PrP^C^ mediates the entrance of tau fibrils in cells, PrP^Sc^ buildup is greatly reduced in their presence, possibly because of an impairment in the prion conversion process.

## Introduction

Tauopathies are a clinically heterogeneous group of diseases that affect many different anatomic regions. Their common trait is the ordered aggregation of misfolded and hyperphosphorylated tau protein within neurons of the central nervous system (CNS). These aggregates have the morphology of fibrils and the tinctorial property of amyloid, i.e. they can be stained by Congo red and thioflavin T. Several *in vitro* and *in vivo* studies showed that, at least to some extent, tau amyloids are able to recruit and seed the aggregation of the endogenous protein, therefore initiating the spreading from neuron to neuron (1-9). These findings, together with the observation that tauopathies progress in the CNS along predictable, neuroanatomically connected circuits (10), led to the hypothesis that they might share some specific features with another group of neurodegenerative diseases, prion disorders. Unlike for prions, tau amyloids do not seem to undergo a fully infectious cycle, and horizontal propagation of tauopathies has not been demonstrated. Hence, tau aggregates are now considered to be “prion-like” proteins, or “prionoids” (11), along with α-synuclein and amyloid-β. Cell-to-cell transfer of tau aggregates is mediated by several mechanisms (endocytosis, receptor-mediated endocytosis, macropinocytosis, tunneling nanotubes) (12-14) that contribute to various extent to the propagation of the amyloids between cells. While some of these pathways are exclusive for tau, some others are exploited also by other neurodegeneration-related proteins. For example, α-synuclein is also thought to be transported along tunneling nanotubes (15).

Recent evidence proposes that the cellular prion protein (PrP^C^) might underlie the progression of several distinct neurodegenerative disorders (16), both promoting the spreading of the misfolded proteins and mediating their neurotoxic effects. Indeed, the presence of PrP^C^ on the cell membrane is essential for the progression of prion disorders, and may be involved in the Aβ-associated neurotoxicity in Alzheimer’s disease (17,18), although the latter finding remains controversial (19). In the case of Parkinson’s disease, PrP^C^ may influence its pathogenesis in two ways: by acting as a receptor for α-synuclein aggregates that facilitates their internalization in healthy cells, and by mediating their toxic effects through the activation of a signaling cascade (20,21). Interestingly, both Aβ- and α-synuclein-associated detrimental effects of PrP^C^, as well as those of bona fide prion diseases, proceed through the activation of the mGluR5 receptor and fyn kinase, corroborating the hypothesis of a common mechanism (18,21,22).

Interestingly, targeting PrP^C^ with antibodies reduces the impairment of long-term potentiation caused by soluble forms of tau, therefore suggesting a role for the prion protein also in the progression of tauopathies (23). Since tau fibrils are mainly intracellular, it is worth investigating if PrP^C^ is involved in the uptake process of the aggregates, as it happens in the case of α-synuclein.

In the present study, we report that PrP^C^ expression enhances the entry of tau fibrils into mouse neuroblastoma cells. Conversely, blocking PrP with antibodies reduced the number of internalized fibrils. These findings suggest that impairing the binding of the fibrils to PrP^C^ might not only to ameliorate synaptic dysfunctions, but also impair the spreading of the aggregates to neighboring neurons. Most surprisingly, we discovered that tau fibrils interfere with PrP^Sc^ accumulation in prion-infected cell lines. We show that the addition of tau K18 amyloids to cultured cells causes a rapid and drastic decrease in PrP^Sc^. We found that the treatment increases the α-cleavage of PrP^C^ into the neuroprotective fragments N1 and C1, thus depleting the pool of misfolding-prone protein and impeding prion conversion.

## Results

### The presence of PrP^C^ on the cell membrane facilitates the uptake of tau K18 amyloids

To test whether PrP^C^ expression affects in some way the entrance of tau K18 amyloids in cells, we compared the number of internalized fibrils in wild-type N2a cells expressing physiological levels of the prion protein to that of N2a cells ablated for PrP^C^ (N2a *Prnp*^-/-^, obtained using the Crispr-Cas9 Based Knockout system (24)). Synthetic amyloids were prepared through an *in vitro* fibrillization process starting from recombinant tau K18 protein (Supplementary Fig. S1 a-d). The purity of the protein was checked by SDS-PAGE (Supplementary Fig. S1a). The resulting amyloids were structurally characterized by atomic force microscopy (AFM) before and after a round of 5 minutes of sonication (Supplementary Fig. S2 a-c). As visible from AFM images, the sonication process both broke the amyloids into more homogeneous smaller species, and disrupts the big aggregates formed as a consequence of the deposition of the sticky fibers in the wells. Short amyloid aggregates were indeed shown to be internalized more easily by cultured cells (25). Therefore, in all following experiments we used exclusively the sonicated preparations of tau K18 amyloids. We performed proteolytic digestion followed by Western blotting, and found that the sonicated preparation of tau fibrils was partially resistant to proteinase K (Supplementary Fig. S1e). We then assessed the cytotoxicity of different concentrations of the fibrils preparations on all the mouse N2a cell lines used in this study for up to 72 hours (Supplementary Fig. S3). We did not detect any significant toxicity after exposure to tau K18 amyloids, which is consistent with the still controversial relationship of the amyloids with neurotoxicity and neuronal degeneration. Only N2a *Prnp*^*-/-*^ showed a small but significant decrease in cell viability upon incubation with tau K18 amyloids; however, the reduction was limited to 14% of total cells and might be related to the increased sensitivity of cells lacking PrP^C^ to a wide range of insults (26,27). To quantify the number of internalized fibrils in N2a cells, we used confocal microscopy of Alexa 488-labelled tau K18 amyloids. Fluorescent labelling of tau K18 amyloids, followed by incubation in cell culture medium and quenching of non-internalized material with the vital dye Trypan Blue (28), allows a more precise quantification of the amyloids by reducing the background signal associated with the use of antibodies. Uptake experiments on wild-type N2a showed that, in our model system and in our experimental conditions, tau K18 amyloids are spontaneously endocytosed in a time- and concentration dependent manner (Supplementary Fig. S4). Higher concentrations (2 μM) required only 24 hours to be taken up by the cells, while in the case of lower concentrations (0.5 μM), 72 hours were needed in order to have 100% of cells positive for aggregates.

To assess the contribution of PrP^C^ to the uptake, we treated N2a and N2a *Prnp*^-/-^ with 2 μM of 488-tau K18 amyloids fibrils for 24 hours, and measured the number of internalized amyloids. We found that wild-type N2a cells took up an average number of 77 ± 41 fibrils per cell compared to the 49 ± 36 fibrils of N2a *Prnp*^-/-^ (p<0.0001, Fig. 1A and B). The difference in the number of internalized tau K18 amyloids between wild-type N2a and N2a *Prnp*^*-/-*^ cells exhibited the same trend in three independent experiments performed with different clones of both cells and 488-tau K18 amyloids. Moreover, we repeated the same experiment varying the concentration of tau K18 amyloids and the incubation time, and found very similar results in terms of differential internalization, therefore confirming the robustness of our findings (Supplementary Fig. S6).

**Figure 1.**
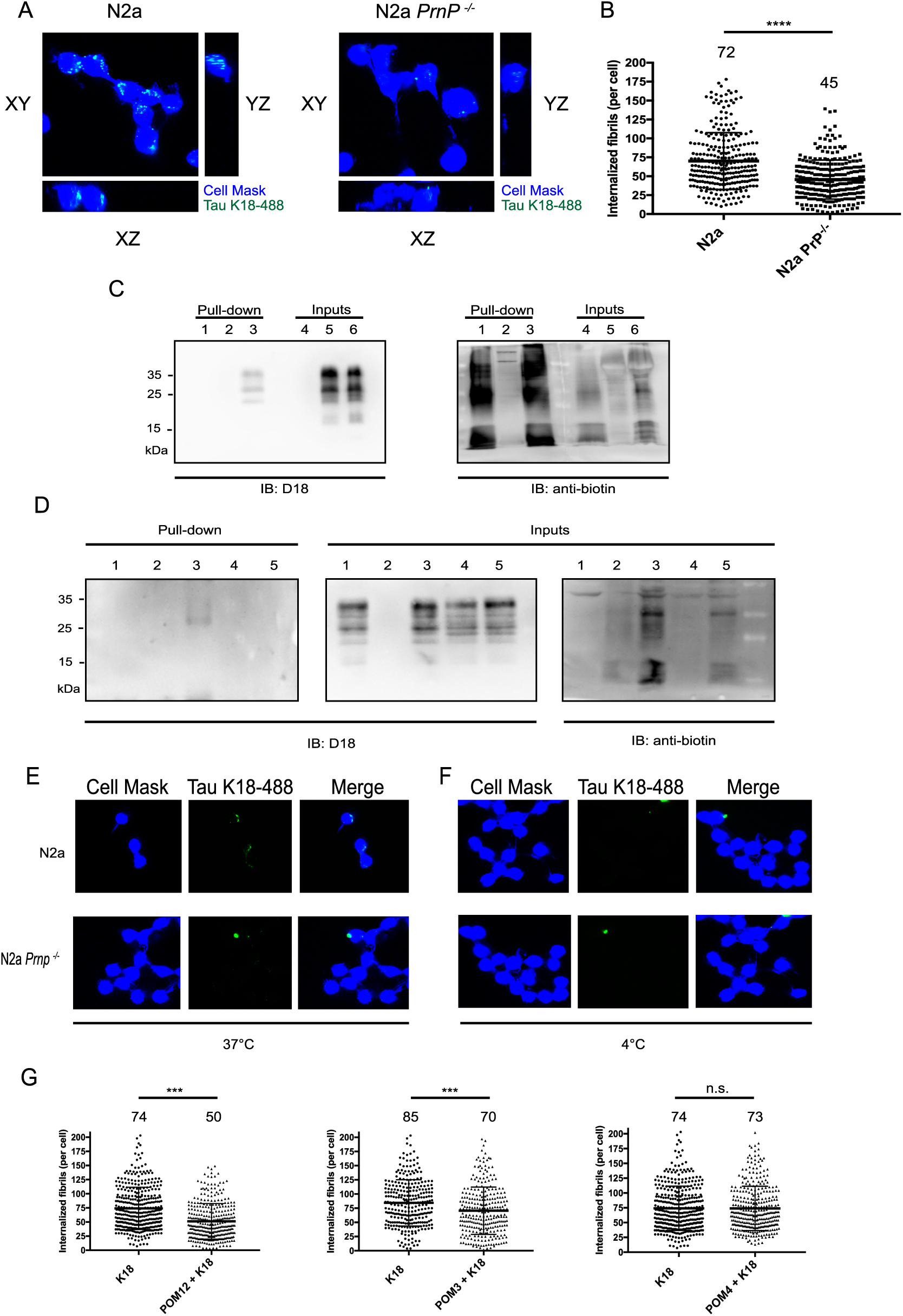
Uptake of tau K18 amyloids in mouse neuroblastoma cell lines. (A) Confocal microscopy images showing the orthogonal views of the central section of the 3D Z-stack for N2a and N2a *Prnp*^-/-^ (in blue) treated with tau K18 fibrils (in green) for 24 hours. White arrows are meant as a help to visualize internalized fibrils. (B) Scatter plot showing the distribution of the number of internalized fibrils in N2a and N2a *Prnp*^-/-^ cells. A total of three hundred cells were counted in blind in three independent experiments. Numbers on top indicate the average number of internalized fibrils. Data were evaluated with unpaired T-test with Welch’s correction. Statistical analysis is indicated as: * = p <0.05, ** = p< 0.01, *** = p<0.001, **** = p<0.0001.. (C) Western Blot of pulled down samples (lanes 1-3) and input samples (lanes 4-6). Lane 1: K18 fibrils only; lane 2: N2a lysate only; lane 3: K18 fibrils and cell lysate. Lane 4: K18 fibrils only; lane 5: N2a cell lysate only; lane 6: K18 fibrils and cell lysate. The membrane was probed with human anti-PrP^C^ antibody D18 and with anti-biotin to visualize tau K18 amyloids. (D) Western Blot of pulled down samples and input samples. Lane 1: N2a lysate only; lane 2: K18 fibrils only; lane 3: K18 fibrils and cell lysate. Lane 4: N2a cell lysate + POM12; lane 5: N2a cell lysate + POM12 + tau K18 amyloids. The membrane was probed with human anti-PrP^C^ antibody D18 and with anti-biotin to visualize tau K18. (E) and (F) Internalization of tau K18 amyloids by N2a and *N2a Prnp*^*-/-*^ cells at 37°C (E) and 4°C (F). Images were acquired using a confocal microscope as series of Z-stacks and the analysis was carried out in 3D. Images show one of the central sections of the stack, both as separate channels and as a merge of the two channels. (G) Scatter plots representing the distribution of internalized fibrils in N2a cells treated only with tau K18 fibrils and in N2a that were pre-treated with different anti-PrP^C^ antibodies. Numbers on top indicate the average number of internalized fibrils. Data were evaluated with unpaired T-test with Welch’s correction. Statistical analysis is indicated as: *= p<0.05; **= p<0.01; ***= p<0.001.

As cells change their volume during the progression of the cell cycle (29), we analyzed the cell dimensions in order to rule out the possibility that the higher internalization of tau K18 amyloids by N2a cells might be artifactual (Supplementary Fig. S5). Indeed, the number of internalized tau K18 amyloids and the cell volume are positively correlated (Supplementary Fig. S5a). No difference was found in the cell volumes between N2a and N2a *Prnp*^*-/-*^ used in the three experiments shown in Fig. 1A and B. (Supplementary, Fig. S5b).

### Tau amyloids bind to PrP^C^ and enter cells through an energy-dependent mechanism

Previous works showed that both Aβ and α-synuclein amyloids bind to PrP^C^ in order to exert their detrimental effects and, in the case of synuclein, to gain cell entrance (17,20). Therefore, we hypothesized that a similar mechanism might underlie also the uptake of tau fibrils. To better define the molecular interaction between the two proteins, we performed a pull-down assay in which biotinylated tau amyloids were used as baits to precipitate PrP^C^ from N2a cell lysate. The data demonstrate that tau K18 amyloids bind to PrP^C^ *in vitro* (Fig. 1C). Evidence of the interaction between the cellular prion protein and monomeric forms of tau is already present in literature (30); however, no previous studies have been conducted using fibrillar aggregates of tau.

As previous reports claim that PrP^C^ interacts with binding partners through its N-terminal tail (20,31), we tried to inhibit the interaction with tau K18 amyloids by pre-incubating N2a cell lysate with POM12 antibody targeting the N-proximal portion (the octapeptide GQPHGGG/SW of the octarepeat region). Lane 5 of Fig. 1D shows that when tau K18 amyloids are incubated with POM12-treated N2a lysate and pulled down, no signal is visible after detection with anti-PrP antibody, confirming that the interaction has indeed been impaired.

The binding of the fibrils to PrP^C^ suggests that the internalization might occur through an energy-dependent, receptor-mediated endocytic pathway rather than via channel diffusion. To test this hypothesis, we incubated N2a and N2a *Prnp*^-/-^ cells with tau fibrils for 6 hours either at 37°C or at 4°C, which blocks all ATP-dependent processes. As expected, wild-type N2a exhibited a greater uptake of tau K18 amyloids under normal culture conditions, with respect to their counterpart not expressing PrP^C^ (Fig. 1E). However, at lower temperature (4°C) we observed an almost complete inhibition of the internalization for both cell types, regardless of their genetic background (Fig. 1F), indicating an ATP-dependent mechanism required for the internalization of tau K18 amyloids.

Finally, we validated the contribution of the cellular prion protein to tau K18 amyloids internalization by targeting PrP^C^ with monoclonal antibodies directed against the N-proximal portion (the aforementioned POM12 antibody), the amino acids 95-100 of the hydrophobic region (POM3 antibody) and a conformational epitope spanning amino acids 121-134 and 218-221 of the C-terminal domain (POM4 antibody)(32). Both POM12 and POM3 decreased the uptake of tau K18 amyloids to a level similar to that of N2a *Prnp*^-/-^cells (Fig. 1G). This is in agreement with the findings of a recent study revealing the toxic effects of soluble tau aggregates are reverted when at least one of these two PrP^C^ regions are blocked (23). On the contrary, targeting the C-terminal domain of PrP^C^ did not result in any change in the amount of internalized tau K18 amyloids (Fig. 1G).

### Tau fibrils induce an increase of the endogenous prion protein that localizes on the cell membrane

To better characterize the mutual relationship between tau amyloids and PrP^C^, we asked if the treatment with tau K18 fibrils has any effect on the endogenous levels of PrP^C^ at different timepoints (Fig. 2, a-b). After 72 hours of incubation, we found an increase of around 60% of the total PrP^C^ level in N2a cells, while at 24 and 48 hours no changes were detected. This effect was dependent on the presence of the aggregates in the culture medium, because when treated cells are passaged and kept in fibrils-free medium, PrP^C^ level went back to basal levels (Fig. 2A). The observed variation in total PrP^C^ occurs only at the protein level, since we did not find any increase in the levels of *Prnp* mRNA after tau K18 amyloids treatment (Supplementary Fig. S7). Immunofluorescence stainings of PrP^C^ in untreated and tau-treated N2a cells revealed that around 35% of the excess PrP^C^ was localized on the plasma membrane (Fig.2, c-d). The increase in PrP-associated fluorescence levels occured only after 72 hours of incubation with tau amyloids, in agreement with Western Blot data (Fig. 2, a-b).

**Figure 2.**
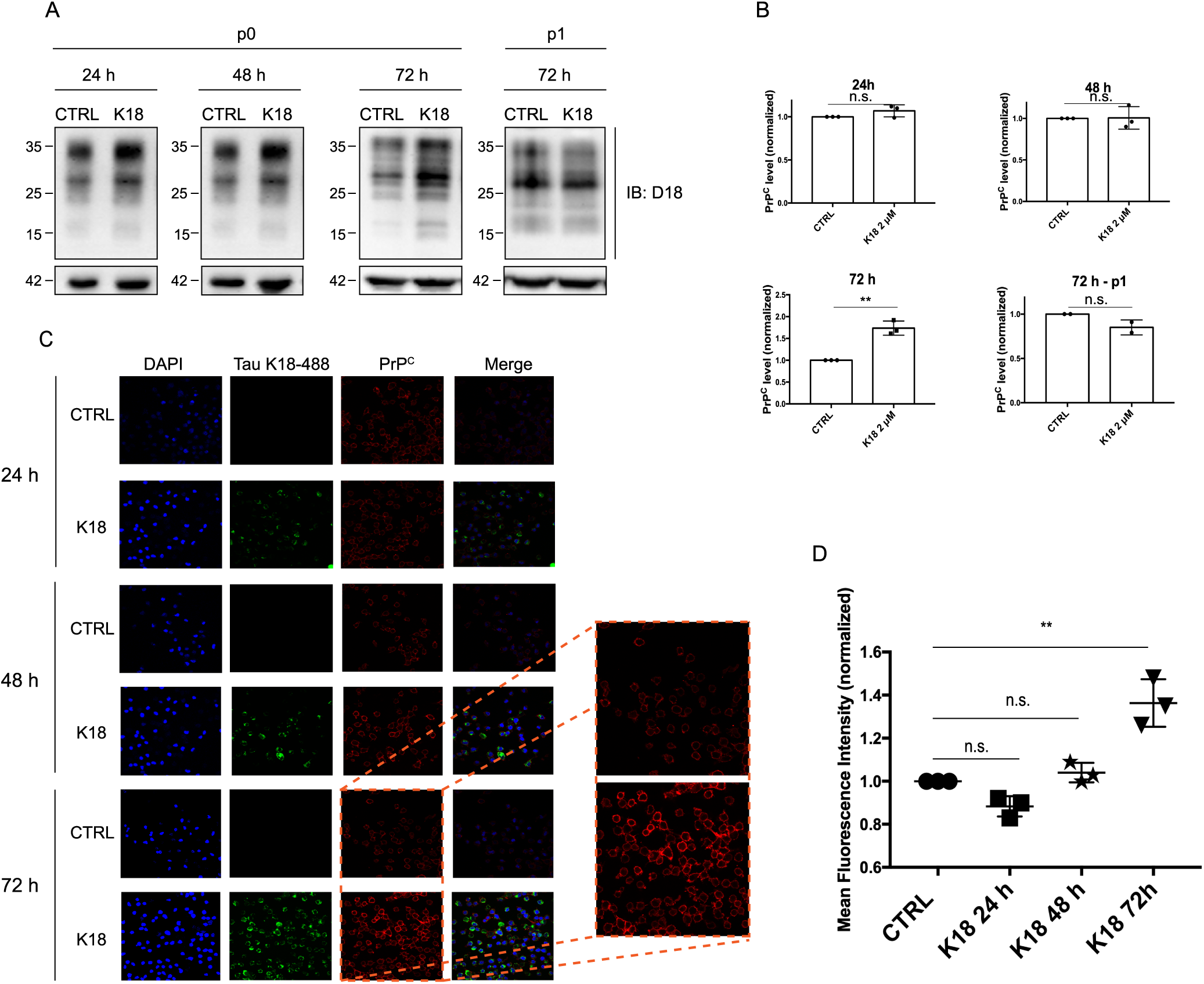
Exposure of N2a cells to tau K18 fibrils induces an increase of the endogenous prion protein. (A) Western blot analysis of N2a cells treated with 2 µM of tau K18 fibrils for 24, 48 and 72 hours and of the first passage of N2a treated for 72 hours. Membranes were probed with human anti-PrP antibody D18. (B) Quantification of three independent experiments. Data are represented as the percentage of total PrP relative to ß-actin. ß-actin is a loading control. Data are represented as mean ± SD. Data were evaluated by unpaired T-test. Statistical analysis is indicated as: n.s.= not significant; *= p<0.05; **= p<0.01; ***= p<0.001. (C) Representative immunofluorescence images of untreated and treated N2a at three different timepoints. Membrane staining for PrP^C^ was performed at 4°C on alive cells using the W226 antibody. The magnified sections show a visible difference in the fluorescence intensity associated to PrP^C^ between control cells and cells treated for 72 hours. The brightness and contrast of some the images has been modified to allow a better visualization. Analysis has been carried out on raw data. (D) Scatter plot of the intensity of the PrP-associated membrane staining of treated N2a compared to control cells. The intensity values, which correlates with the levels of PrP on the membrane, is reported as the ratio between the average intensities of treated and control cells for each timepoint. Controls have been normalized to 1. The three values represent three independent experiments, in which 700 cells for each condition were analysed. Data have been evaluated with unpaired T-test. Statistical analysis is indicated as: n.s.= not significant; *= p<0.05; **= p<0.01; ***= p<0.001.

### Tau amyloids lower PrP^Sc^ levels in cell lines infected with different prion strains

Although the process of prion conversion and PrP^Sc^ formation depends mainly on the presence of PrP^C^, recent studies revealed that it can be affected by the concomitant presence of intracellular amyloids of other neurodegeneration-related proteins (20). In the case of tau protein, this assumes clinical relevance as some cases were documented where prion pathology co-exists with tau deposits (33-35). To assess the effect of tau fibrils on PrP^Sc^ levels, we exposed a neuroblastoma cell line persistently infected with RML prion strain (ScN2a RML) to two different concentrations of tau K18 amyloids and we analysed the PK-resistant PrP^Sc^ at different timepoints (Fig. 3A). The quantification of three independent experiments (Fig. 3B), showed that the time required for PrP^Sc^ clearance is dependent on the concentration of tau K18 amyloids. Indeed, cells treated with 5 µM tau K18 amyloids showed significantly less prions after 48 hours, while treating with a lower concentration of tau K18 amyloids (2 µM) required 72 hours to lead to a 50% decrease in prion burden. Although the highest concentration of tau K18 amyloid proved to be more effective in clearing PrP^Sc^ after 72 hours, we chose to use 2 µM for further experiments, as it gave more reproducible results. Immunofluorescence data (Fig. 3C) and Western Blot analysis (Supplementary) revealed that tau K18 amyloids are taken up by ScN2a cells, and they appear to co-localize with lysosomes (Supplementary Fig. S8). Next, we wondered if PrP^Sc^ levels could remain low upon passaging, even in the absence of tau fibrils. After a 72 hours-incubation with 2 µM of tau K18 amyloids, cells were kept in culture for 5 additional passages. (Fig. 4A, 4B, 4C). The prion load decreased after the treatment with the amyloids and remained low through two subsequent passages (Fig. 4A and B), while monomeric K18 used as a control did not affect prion levels. However, although the reduction observed after the initial passages was quite pronounced, it was not maintained after the third passage without fibrils, where PrP^Sc^ levels rose again to their initial level (Fig. 4C). A longer incubation with the fibrils did not result in a complete clearance of PrP^Sc^, since a weak signal was still present at passage 3 (Supplementary, figure S9).

**Figure 3.**
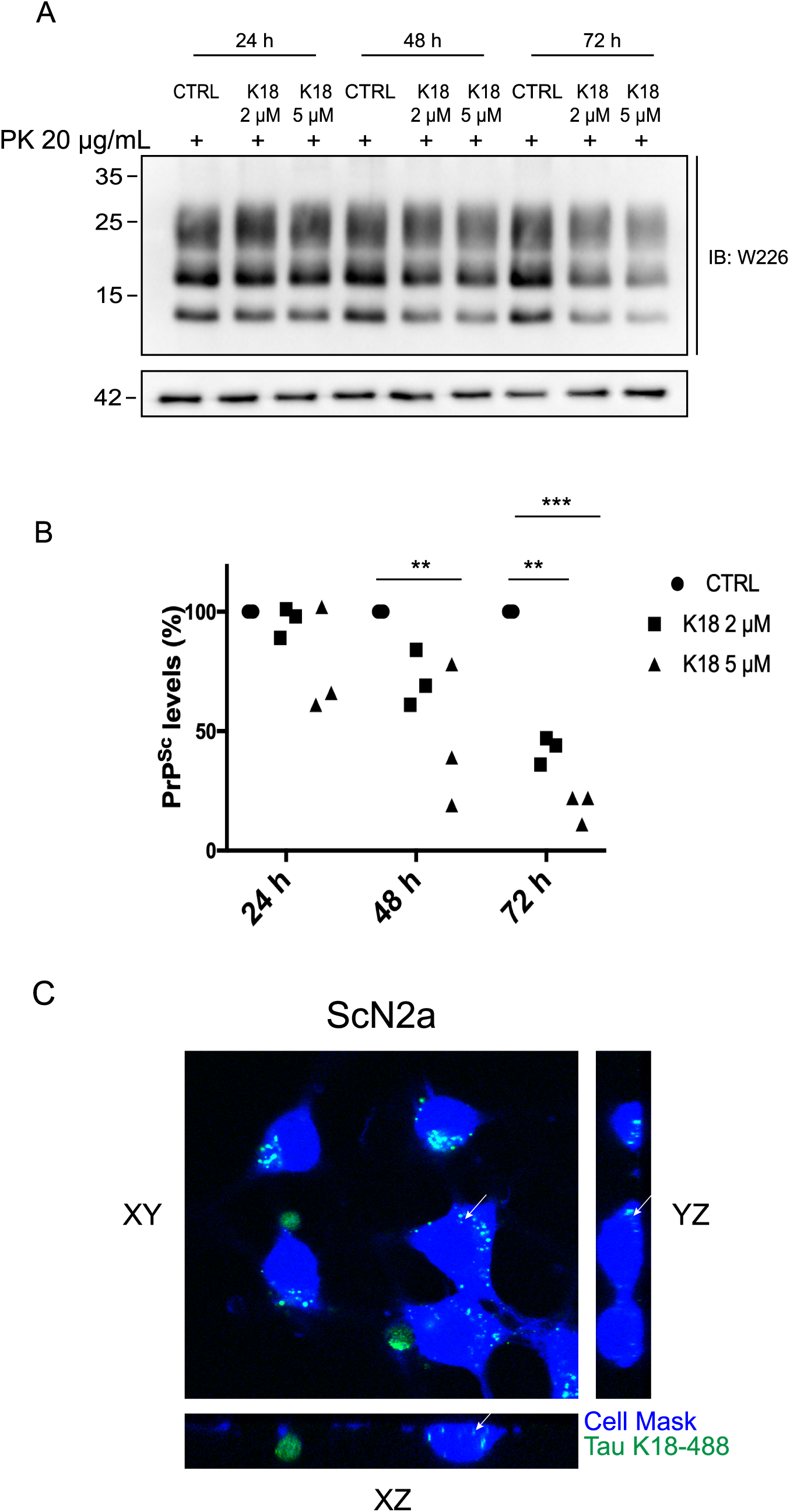
Tau-induced PrP^Sc^ clearance in ScN2a RML cell line at different timepoints. (A) Western Blot analysis of PK-resistant PrP^Sc^ upon treatment with different concentrations of tau K18 amyloids at 24 hours, 48 hours and 72 hours. (B) Quantification of three independent experiments. Values are shown as a percentage of PK-resistant form relative to β-actin. β-Actin is a loading control. Data are represented as mean ± SD. Data were evaluated by one-way ANOVA with multiple comparisons. Statistical analysis is indicated as: *= p < 0.05, **= p < 0.01, ***= p < 0.001. (C) Confocal microscopy image showing the orthogonal views of the central section of the 3D Z-stack of ScN2a RML cells treated with 2 µM of tau K18 amyloids (in green) for 72 hours.

**Figure 4.**
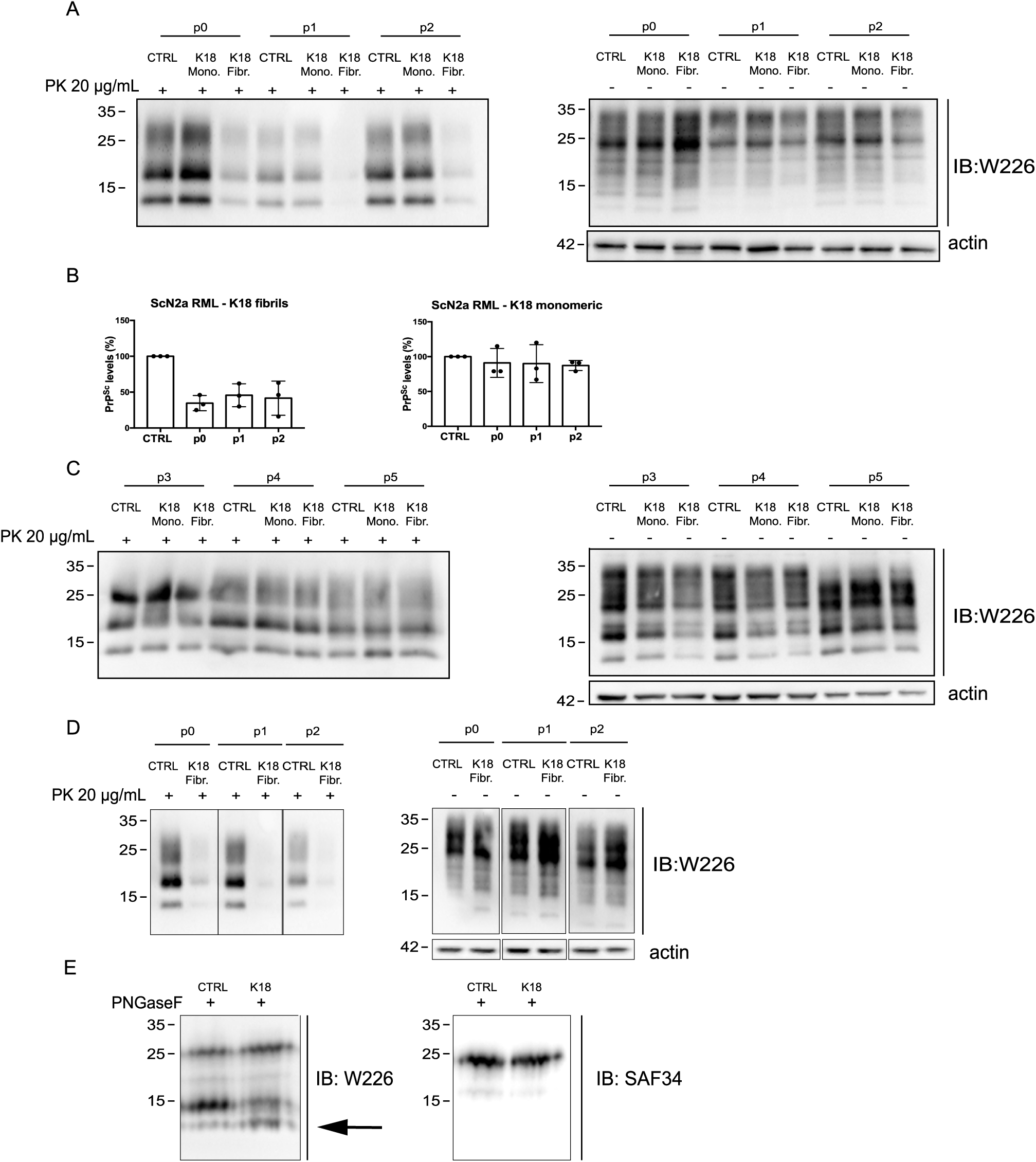
Clearance of PrP^Sc^ from prion-infected ScN2a cell lines after treatment with tau K18 amyloids. (A) Western blot analysis of PK-resistant PrP^Sc^ in ScN2a RML cell lysates upon treatment with tau K18 amyloids for 3 days from the addition of the fibrils (passage 0) to passage 2. (B) Quantification of three independent experiments. Values are shown as a percentage of PK-resistant form relative to β-actin. β-Actin is a loading control. Data are represented as mean ± SD. Data were evaluated by one-way ANOVA with multiple comparisons. Statistical analysis is indicated as: *= p < 0.05, **= p < 0.01, ***= p < 0.001. (C) Western Blot analysis of further passaging of treated cells shown in (A), from passage 3 to passage 5. (D) Western blot analysis of PK-resistant PrP^Sc^ in ScN2a 22L cell lysates upon treatment with tau K18 amyloids for 3 days. (E) Electrophoretic pattern of PNGase F digested PrP in ScN2a control cells and cells treated with tau K18 fibrils. Solid black arrow shows the presence of C1 fragment recognized with C-terminal Ab (W226), while the broken black arrow indicates the absence of the bands when the membrane was probed with N-terminal antibody (SAF34).

We tested the strain dependency of tau K18-induced clearance of PrP^Sc^ by repeating the treatment on ScN2a infected with the 22L prion strain (Figure 3C). Also in this case we reported a strong reduction in the PrP^Sc^ load after the administration of the fibrils, indicating that the mechanism activated by tau K18 is not affected by strain differences.

Taken together, these results show that tau K18 amyloids are able to clear PrP^Sc^ in cultured cells permanently replicating different prion strains, and that they have to be present in the cell culture medium in order to keep the prion burden low. This effect occurs probably through a direct interaction with either PrP^C^ or PrP^Sc^. Since tau K18 amyloids interact with PrP^C^, we hypothesized that the fibrils might bind and reduce the availability of PrP^C^ for PrP^Sc^-induced conversion. Indeed, we found that the treatment with K18 fibrils promotes the α-cleavage of PrP^C^ with the formation of the N1 and C1 fragment, thus reducing the pool of PrP^C^ needed for the conversion (Fig. 4E).

## Discussion

A number of reports claim that the role of the cellular prion protein in neurodegeneration is not limited to prion disorders, but extends also to the pathogenesis of Alzheimer’s disease and synucleinopathies(16,17,20,23,36). Indeed, several reports indicate that PrP^C^ mediates the neurotoxicity of several protein aggregates (α-synuclein, amyloid β) in two different ways. Firstly, PrP^C^ was reported to act as a membrane receptor to facilitate their internalization(20,23,37,38). Secondly, PrP^C^ may function as a transducer of their detrimental effects through the activation of metabotropic glutamate receptors(18,39).

Regarding tau, the impairment in long-term potentiation caused by soluble tau oligomers might be ameliorated by directly targeting PrP^C^ in affected animals (23). However, the exact mechanism by which PrP^C^ contributes to tau pathology remains unknown. In this study, we investigated whether the expression of PrP^C^ might affect the internalization of tau amyloids. First, we showed that sonicated tau K18 fibrils bind to PrP^C^. The existence of an interaction between the two partners is the first mandatory step in the cascade of events leading to PrP^C^-dependent internalization. Although there is proof of the interaction of the prion protein with monomeric full-length tau in the literature (30), no experiments had been performed so far using synthetic aggregates. We found that cells that express PrP^C^ on the cell surface (N2a) internalize more tau amyloids compared to the same cells from which the gene encoding PrP^C^ was removed (N2a *Prnp*^-/-^). N2a *Prnp*^-/-^ cells, even if deprived of PrP^C^, were able to internalize a discrete number of fibrils, consistently with previous reports of multiple mechanisms occurring at the same time and independently from each other. However, when the prion protein was present, the number of internalized fibrils almost doubled, meaning that the PrP^C^-mediated pathway contributed strongly to this process. The role of PrP^C^ was orthogonally confirmed by the observation that the targeting of the N-terminus or the hydrophobic domain of PrP^C^ using antibodies lead to a reduction of the internalized fibrils. Most intriguingly, we discovered that the interaction between PrP^C^ and tau fibrils has mutual effects on both partners; while the former acts as a receptor, the presence of tau aggregates leads to an increase of the cellular level of the prion protein, which is re-localized mainly on the plasma membrane where it could perform its receptor activity. The increase in PrP^C^ could be due either to the trafficking of the newly synthesized PrP^C^ on the surface, or to a longer retention on the membrane as a consequence of the interaction with the fibrils, as no changes in *Prnp* expression levels were detected. Given that the prion protein facilitates the internalization of tau amyloids, we deem it likely that a higher amount of PrP^C^ exposed outside the cells might result in more extracellular tau fibrils gaining cell entrance Although this phenomenon might be a simple side-effect of the interaction and subsequent internalization of the aggregates, we deem it likely that tau fibrils are boosting their own spreading by increasing the number of receptor molecules on the cell surface. A similar hypothesis has been proved true in the case of intercellular transfer of tau amyloids through tunneling nanotubes, where extracellular tau species (monomers and fibrils) activated the formation of new nanotubes and therefore facilitated fibrillar tau transport between neurons (15). On the other hand, protein aggregates (tau and synuclein) are trafficked in early endosomes and lysosomes after the internalization (20,40,41). The need of the cell to deal quickly with a large amount of exogenous and potentially dangerous material might clog the degradation system, with subsequent impact on the normal turnover of the endogenous proteins. Indeed, a similar effect on PrP levels has been reported after treatment with synuclein fibrils (20), although the localization of the excess protein was not investigated. Alternatively, the binding of the amyloids to PrP^C^ on the cell membrane might simply impede its internalization and retain the protein in its position, therefore leading to a general increase as the newly synthesized protein follows its normal pathway all the way up to the surface.

Moreover, we observed that the scrapie load in cell cultures decreased after incubation with tau K18 fibrils. Tau-mediated clearance was independent of the PrP^Sc^ conformation, as a similar reduction was observed for both RML and 22L prion strains. One way in which tau K18 amyloids might lead to PrP^Sc^ reduction is through a direct binding to either PrP^C^ or PrP^Sc^, inhibiting prion conversion by depleting respectively the pool of the substrate or of the template. Indeed, tau K18 fibrils bind to PrP^C^ and promote its processing into the N1 and C1 fragments, which reduces the amount of cellular prion protein available for the conversion. The similar effects observed after the exposure of two different prion strains to tau aggregates seems to suggest that PrP^Sc^ might not be the target. As the exact structure of PrP^Sc^ is undefined, the structural differences distinguishing the RML and 22L prion strains are not known, and could be so subtle not to impact the binding of K18 fibrils in a detectable manner. In view of the wealth of data suggesting that PrP^C^ interacts with many amyloid, the most parsimonious interpretation is that tau K18 amyloids interacts with the cellular prion protein PrP^C^. In conclusion, here we provide evidence of another relevant role for PrP^C^ in mediating the internalization of exogenous tau fibrils, therefore contributing to the cell-to-cell spreading of tau K18 amyloids and to the onset of the pathology. The interaction between tau fibrils and the cellular prion protein seems to affect also the process of prion replication in prion-infected cell lines. This occurs probably through the inhibition of the mechanism of PrP^C^ conversion into PrP^Sc^, due to an increased rate of α-cleavage of PrP^C^ stimulated by the binding to tau fibrils.

## Materials and methods

### Production and fibrillization of recombinant tau K18

Expression and purification of recombinant tau K18 fragment were performed as previously described (42). *In vitro* fibrillization reactions were prepared in a 96-well black plate with transparent bottom (BD Falcon) in a final volume of 200 µL per well. Fibrillization reaction was composed as follows: tauK18 0.5 mg/mL, heparin 10 µg/mL, DTT 0.1 mM, PBS 1X pH 7.4. Due to its toxicity to cultured cells, 10 µM Thioflavin T was added only in 4 wells that were used to monitor the aggregation reaction in real-time. Each well contained also a 3-mm glass bead (Sigma). The plate was covered with sealing tape (Fisher Scientific) and incubated at 37°C under orbital shaking (50 seconds of shaking at 400 rpm followed by 10 seconds of rest) on FLUOstar Omega (BMG Labtech) microplate reader. Fluorescence was monitored every 30 minutes by bottom reading at 444 nm of excitation and 485 nm of emission. For cell culture experiments, fibrillization was perfomed as described above but in absence of ThT. The reaction was stopped after 15 hours, when fluorescence reached plateau. Newly formed aggregates were pelleted by ultracentrifugation (55000 rpm for 1 hour at 4°C), resuspended in an equal volume of sterile PBS and stored at −20°C. Before use, aliquots were thawed and sonicated for 5 minutes in an ultrasonic bath (Branson 2510).

### Cell culture

Mouse neuroblastoma cells N2a were kindly provided by Prof. Chiara Zurzolo (Unité de traffic membranaire et pathogenèse, Institute Pasteur, Paris, France). N2a *Prnp*^-/-^ cells were kindly provided by professor Gerold Schmitt-Ulms (Tanz Centre for Research in Neurodegenerative Diseases, University of Toronto, Toronto, Ontario, Canada), for which they used the CRISPR-Cas9-Based Knockout system to ablate the expression of PrP protein [293]. ScN2a cells are clones persistently infected with the RML prion strain as described by Prusiner’s group [437] or with the 22L prion strain. Cells were grown at 37°C and 5% CO2, in minimal essential medium (MEM) + glutamax (Thermo Fisher Scientific Inc.), supplemented with 10% fetal bovine serum, 1% non-essential aminoacids, and 100 units/ml penicillin and 100 μg/ml streptomycin.

### TauK18 fibrils infection in cell lines

TauK18 amyloids were added to the cell culture medium of N2a, N2a *Prnp*^-/-^ and ScN2a cell lines in 10 cm plates, and incubated for a variable amount of time according to the specific experimental setting (ranging from 1 day to 6 days). For the evaluation of PrP^Sc^ clearance in ScN2a cell line, cells were split and maintained for five additional passage in fibrils-free cell culture medium. For fluorescence quantification of amyloids internalization, cells were plated in 24-wells plates on 12 mm coverslips and incubated with Alexa488-labeled fibrils for 24 – 72 hours.

### Pull-down assay

N2a cells were lysed in lysis buffer (0.1% NP40, 50 mM TrisHCl pH 8, 150 mM NaCl) containing one tablet of Complete™ ULTRA Tablets, EDTA-free, glass vials Protease Inhibitor Cocktail (Roche). Total protein content was quantified using bicinchoninic acid protein (BCA) quantification kit (Pierce). For inhibition of the interaction using POM12, N2a cell lysate was incubated for 1 hour with POM12 (15 µg/mL) at 37°C. 1 mg of total protein content was diluted in lysis buffer to a final volume of 500 µL, then biotinylated tauK18 fibrils were added to a final concentration of 2 µM. Samples were left standing on a rotor wheel overnight at 4°C. The day after, 30 µL were collected from each sample and stored at −20°C for further analysis (input samples). 30 µL of neutravidin bead slurry (NeutrAvidin Agarose Resins, ThermoFisher Scientific) were added to each tube, and the samples were incubated on the rotor wheel at 4°C for 4 hours. To precipitate the beads, the samples were centrifuged at 4°C for 2 minutes at 2000 rpm, the pellet was washed with three subsequent steps of resuspension in lysis buffer and centrifugation, and finally resuspended in 20 µL of Laemmli loading buffer. Samples were boiled for 10 minutes and centrifuged at 13000 rpm for 1 minutes to detach and pellet the beads, respectively. The supernatants were collected and stored at −20°C for further analysis.

### Trypan blue quenching of non-internalized fibrils and imaging

30000 cells were cultured on each coverslip and treated with different concentrations of Alexa488-labeled tau K18 amyloids. Before fixation, cells were washed twice with sterile PBS and incubated for 5 minutes with a sterile 1:1 solution of Trypan Blue : PBS. As shown by Karpowicz et al.(28), Trypan Blue quenches green fluorescence through an energy-transfer mechanism, but being unable to enter alive cells, only the fluorescence coming from non-internalized fibrils is quenched.

Cells were then rinsed three times with sterile PBS and fixed for 10 minutes with 4% paraformaldehyde/PBS. After three washing steps, cells were permeabilized for 5 minutes with 0.2% Triton X-100/PBS, rinsed again with PBS and incubated for 1 hour with HCS Blue Cell Mask (ThermoFisher Scientific) diluted 1:1000, a specific dye that labels the whole cell cytoplasm. Coverslips were mounted in Fluoromount-GTM (ThermoFisher Scientific) and stored at 4°C for confocal fluorescence microscopy. Images were acquired using a Nikon confocal microscope (Nikon C1).

### Inhibition of tau K18 amyloids internalization using POM monoclonals

Anti-PrP^C^ monoclonal antibodies were kindly provided by Prof. Adriano Aguzzi (Institute of Neuropathology, University of Zürich) (32). N2a cells plated on coverslips were pre-treated with each POM antibody (15 µg/mL) for 1 hour at 37°C, followed by incubation with Alexa488-labeled tauK18 fibrils for 24 hours. Coverslips were processed and mounted as described above. To check the binding of POM antibodies to membrane PrP^C^, cells treated with POM antibodies for 1 hour were fixed with 4% paraformaldehyde:PBS for 10 minutes, then blocked with 2% FBS for 30 minutes and incubated with Alexa594-conjugated goat anti-mouse IgG diluted 1:200 for 1 hour.

### Uptake quantification

The uptake quantification was performed in blind on a total of four hundred cells in three independent experiments. Random fields on each coverslip were captured at 63x magnification (digital zoom 3X) using Nikon C1 confocal microscope. Images were acquired as stacks of 30 −40 optical sections of 0.27 µm, 526 x 526, and subsequently deconvolved with a 3D deconvolution algorithm (in blind) of the NIS Elements software. Deconvolution was applied only to the green channel, as it allows a better separation of the green dots representing the fibrils. Deconvolved images were analysed using Volocity Workstation (PerkinElmer, version 4.1) and internalized fibrils were counted using a home-made script which intersects the objects recognized in the green channel with the 3D reconstruction of the cellular volume (blue channel). To discriminate the signals corresponding to fibrils from the background, we used the volume of a single monomer of synuclein as a threshold. Synuclein is very similar to tau K18 in terms of molecular weight and unfolded structure. The volume of a monomer in its multiple conformations was obtained from Zhang et al. (43)

### Membrane immunostaining of PrP^C^

N2a cells seeded on coverslips were treated for 24 – 72 hours with 2 µM of tau K18 amyloids. Surface staining of PrP^C^ was performed as described in Stincardini et al. (44). Cells were incubated with W226 antibody diluted 1:250 in Opti-MEM (Life Technologies) for 15 minutes at 4°C, followed by washing in PBS and fixation with 4% paraformaldehyde:PBS for 10 minutes. Coverslips were then washed three times with PBS, incubated with blocking solution (2% FBS in PBS) for 30 minutes and incubated with Alexa-594 conjugated goat anti-mouse IgG (Invitrogen) diluted 1:200 in blocking solution. Finally, nuclei were counterstained with DAPI (1:1000). Coverslips were mounted in Fluoromount-GTM (ThermoFisher Scientific) and stored at 4°C. Images were acquired with Nikon C1 confocal at 63x magnification, both as 2D-images of medial planes and series of 30-40 optical sections (z-step = 0.27 µm, 526×526).

### Quantification of membrane staining

Quantification of membrane staining was performed with Volocity Workstaton (PerkinElmer) on 2D images. Image segmentation consisted first of nuclei identification by the DAPI signal, and selection of the region of interest by the Alexa-594 signal. Mean fluorescence intensity values of Alexa594-conjugated anti-PrP antibody were measured for more than 600 objects per coverslip, and weighted averages were calculated in order to identify any difference between untreated and treated samples. To confirm that 2D quantification is representative of the situation of the whole cell membrane, the same 3D analysis was performed on a number of z-stacks from both untreated and treated samples.

### Western blotting

Total protein contents of N2a untreated and treated with 2 µM of tau K18 amyloids were quantified using bicinchoninic acid protein (BCA) quantification kit (Pierce). Fifty µg of total proteins were used and additioned with 5X loading buffer in a ratio 1:5. The samples were boiled at 100°C for 10 minutes, loaded onto a 12% Tris-Glycine SDS-PAGE gel and transferred onto Immobilon P PVDF membranes (Millipore) for 2 hours at 4°C. The membrane was blocked in 5% non-fat milk and incubated overnight at 4°C with human anti-PrP antibodies (D18 1:1000, W226 1:1000, SAF34 1:1000). The membrane was washed three times with TBS buffer additioned with 0.2% Triton X-100, incubated for 45 minutes at room temperature with goat anti-human IgG conjugated with horseradish peroxidase and developed with the enhanced chemiluminescent system (ECL, Amersham Biosciences). After the acquisition, the membrane was incubated for 30 minutes at room temperature with anti β-actin (1:10000, A3854 Sigma-Aldrich), wich is used as a normalizer. Images were acquired with Uvitec Alliance (Cambridge) and densitometric analysis was performed using Uviband analysis software. Data are expressed as mean ± SD, and the values of the controls are adjusted to 100%. For tau fibrils detection, the membrane was probed with anti-4R tau RD4 antibody diluted 1:1000.

### PNGase F treatment

PNGase F treatment (New England Biolabs) was performed on control and K18-treated ScN2a cell lysates. Twenty µg of total proteins were incubated with 1 µL of Glycoprotein Denaturating Buffer (10X) and water to reach a total reaction volume of 10 µL, then the mix was heated at 100 °C for 10 minutes to denature the proteins. The solution was chilled on ice and centrifuged for 10 seconds, then additioned with 2 µl of GlycoBuffer 2 (10X), 2 µl of 10% NP-40, 6 µl of H2O and 1 µl of PNGase F. The solution was incubated overnight at 37°C. Deglycosylated proteins were then analysed by Western Blot.

## Acknowledgements

Funding was provided by JPND Reframe Consortium to G.L.

E.D.C. would like to acknowledge Fabio Perissinotto, PhD, for the AFM analysis of tau K18 amyloids, Rabah Soliymani, PhD, for Mass Spectrometry analysis and Camila Gherardelli for help and support with the experiments of tau amyloids internalization.

## Author contributions

E.D.C. conceived and performed the experiments, wrote the manuscript. L.C. Performed experiments of PrP^Sc^ clearance by tau K18 amyloids. S.V. Performed pull-down experiments. M.G. Support with microscopy images acquisition and analysis. A.A. Suggested experiments and revised the paper. G.L. Conceived and designed the experiments.

## Footnotes

## The abbreviations used are

PrP^C^: cellular prion protein
PrP^Sc^: scrapie prion protein
mGluR5: metabotropic glutamate receptor 5
N2a: murine neuroblastoma cell line
AFM: Atomic Force Microscopy
PK: proteinase K

