## Supplementary Files for "The uptake of tau amyloid fibrils is facilitated by the cellular prion protein and hampers prion propagation in cultured cells"

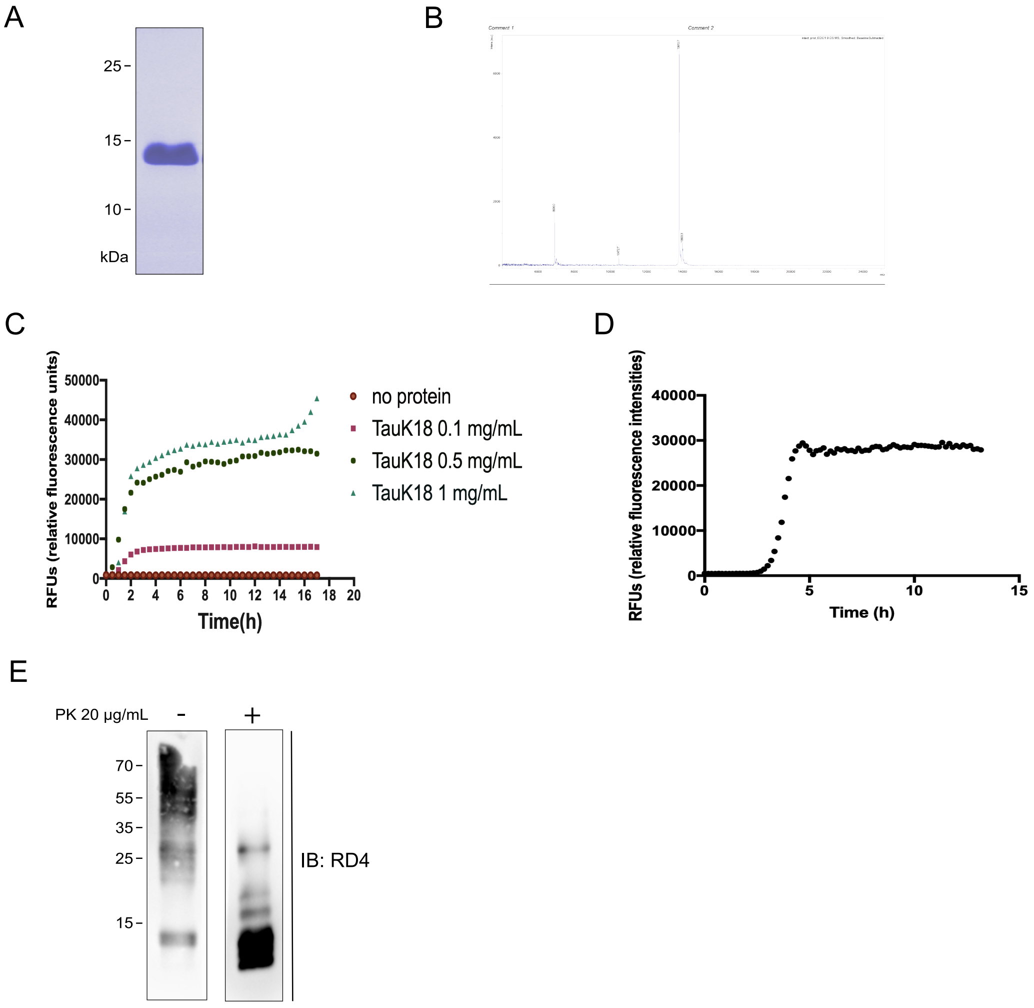


**Supplementary Figure S1. Expression and purification of recombinant tau K18 protein.** (A) Purified K18 fragment of tau protein. (B) Mass spectrometry confirmed the molecular mass of tau K18 protein. (C) Fibrillization curves of tau K18 at different concentrations monitored by ThT fluorescence. A condition with no protein was added as negative control. (D) Fibrillization curve of tau K18 at 0.5 mg/mL of concentration. Black arrow indicates the time point for the collection of tau fibrils. (E) Western blot depicts the biochemical profile of tau K18 fibrils after 5 minutes of sonication, before and after proteinase K digestion.

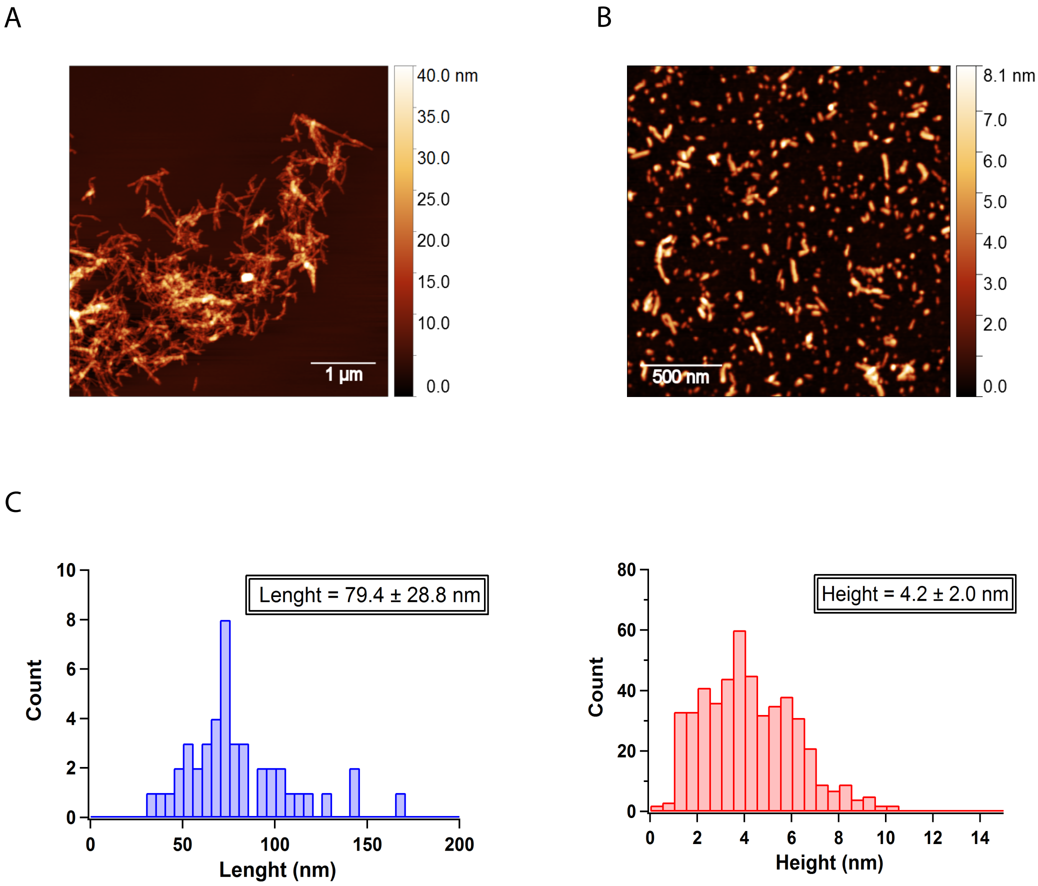


**Supplementary Figure S2. AFM analysis of tau K18 fibrils.** AFM images of non-sonicated (A) and sonicated (B) tau K18 fibrils. (C) Length and height quantification of sonicated tau K18 fibrils.

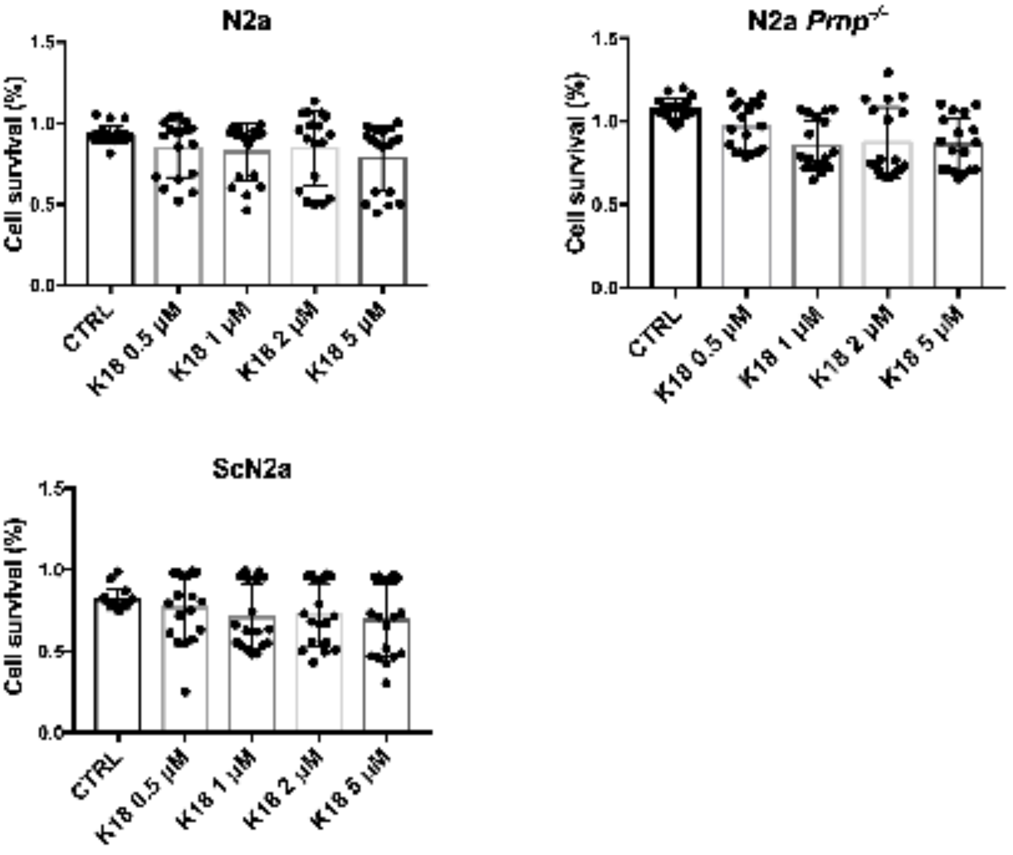


**Supplementary Figure S3. In vitro cell cytotoxicity.** Effect of different concentrations (0.5 µM, 1 µM, 2 µM, 5 µM) of sonicated tau K18 fibrils on the viability of N2a $PrP^{+/+}$, N2a $PrP^{-/-}$ and ScN2a measured by MTT assay. Data represent three separate experiments, each performed in six replicates.

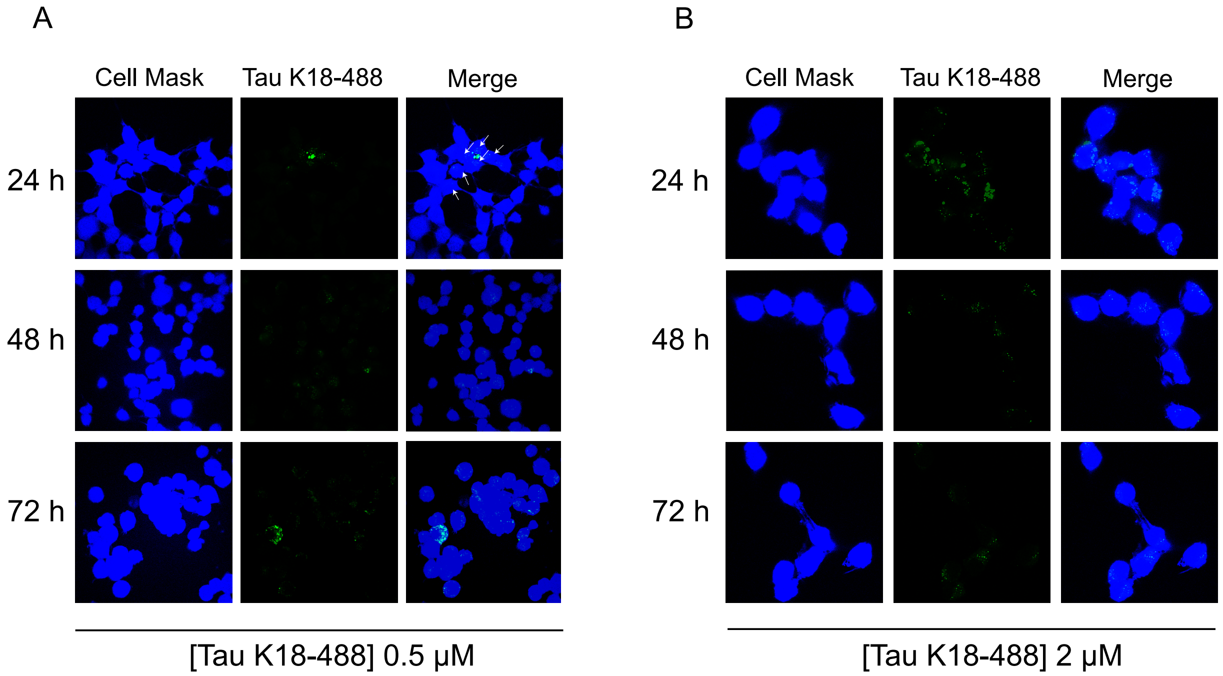


**Supplementary Figure S4. Internalization of tau K18-488 fibrils in N2a cells at different timepoints.** N2a cells were incubated with (A) 0.5 µM or (B) 2 µM of tau K18-488 fibrils for 24, 48 and 72 hours. Images were acquired as series of Z-stacks and the analysis was carried out in 3D. Panels A and B show the XY projection of one of the central sections of the stack. White arrows in panel A indicate the cells positive for tau aggregates.

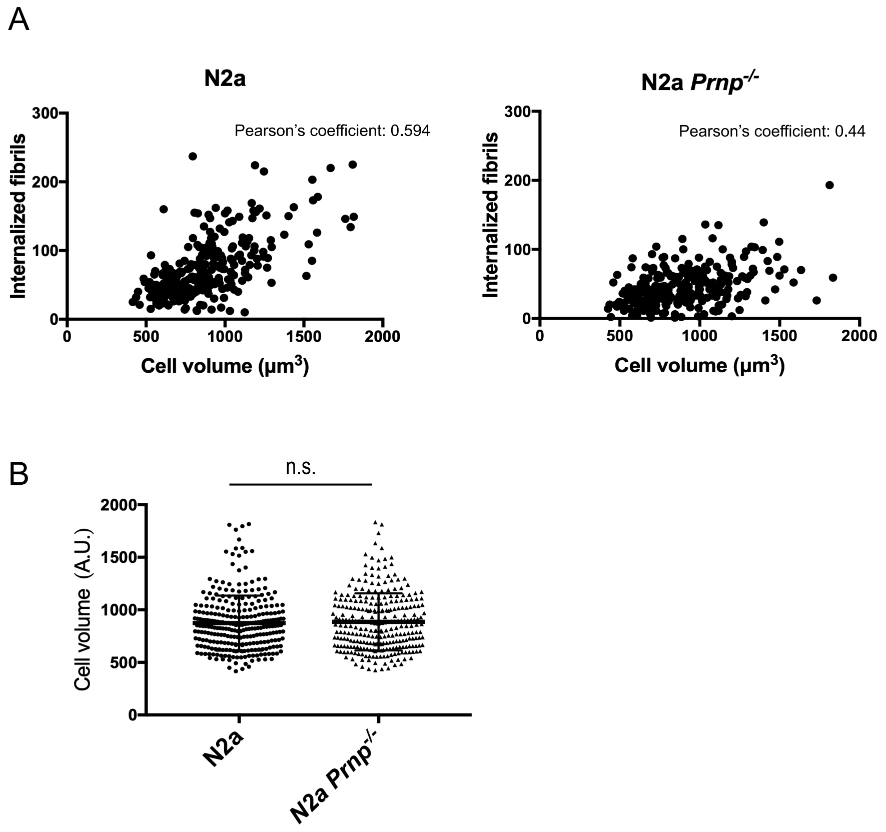


**Supplementary Figure S5. Correlation between cell volume and the number of internalized fibrils per cell in N2a and N2a KO cell lines.** For both cell types a direct correlation exists, and can be defined as "moderate correlation" according to Pearson's coefficient.

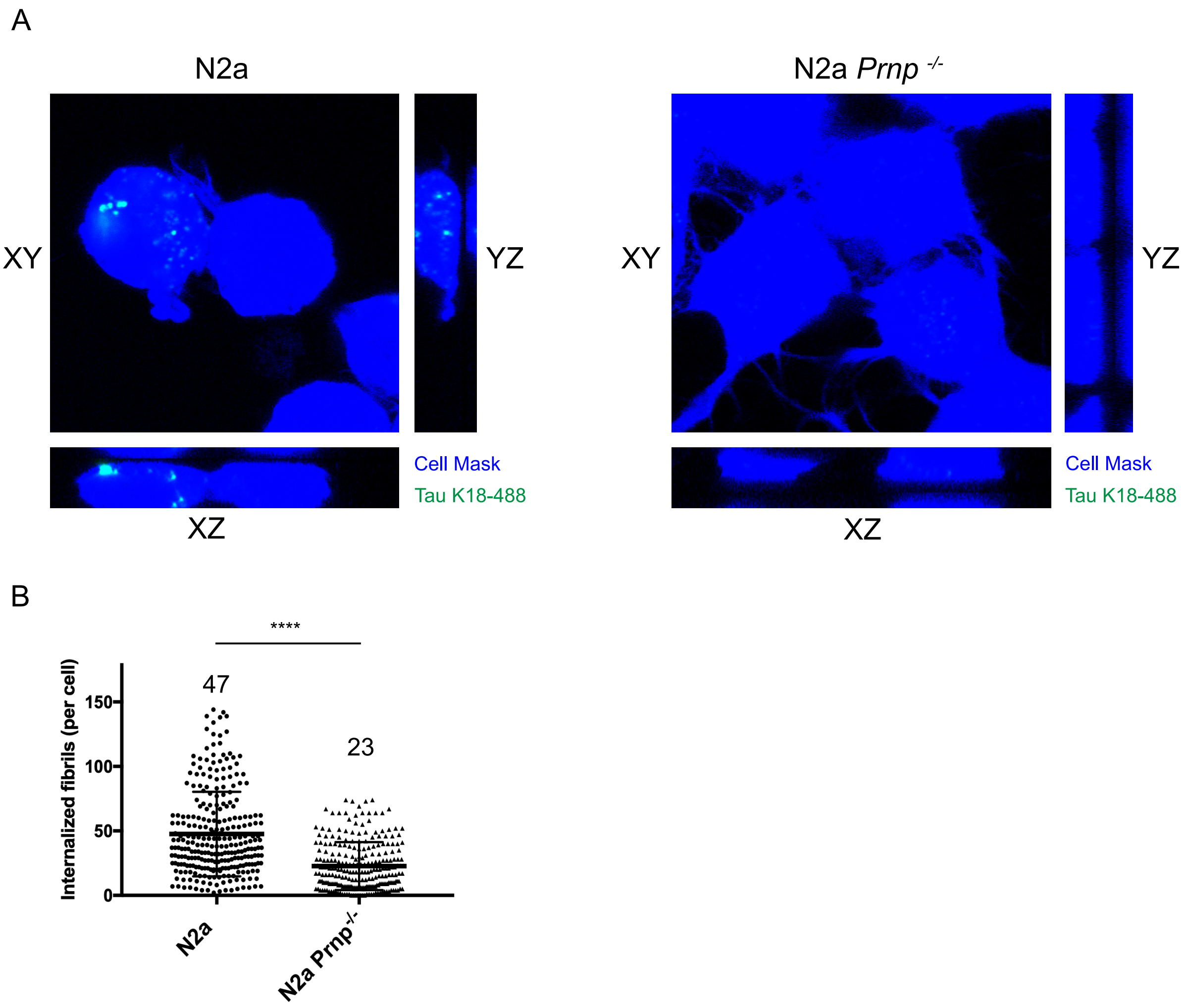


**Supplementary Figure S6. Uptake of tau K18 amyloids in mouse neuroblastoma cell lines.** (A) Confocal microscopy images showing the orthogonal views of the central section of the 3D Z-stack for N2a and N2a *Prnp*$^{-/-}$ (in blue) treated with 0.5 μM tau K18 fibrils (in green) for 72 hours. (B) Scatter plot showing the distribution of the number of internalized fibrils in N2a and N2a *Prnp*$^{-/-}$ cells. A total of three hundred cells were counted in blind in three independent experiments. Number on top indicate the average number of internalized fibrils per cell. Data were evaluated with unpaired T-test with Welch's correction. Statistical analysis is indicated as: * = p <0.05, ** = p< 0.01, *** = p<0.001, **** = p<0.0001.

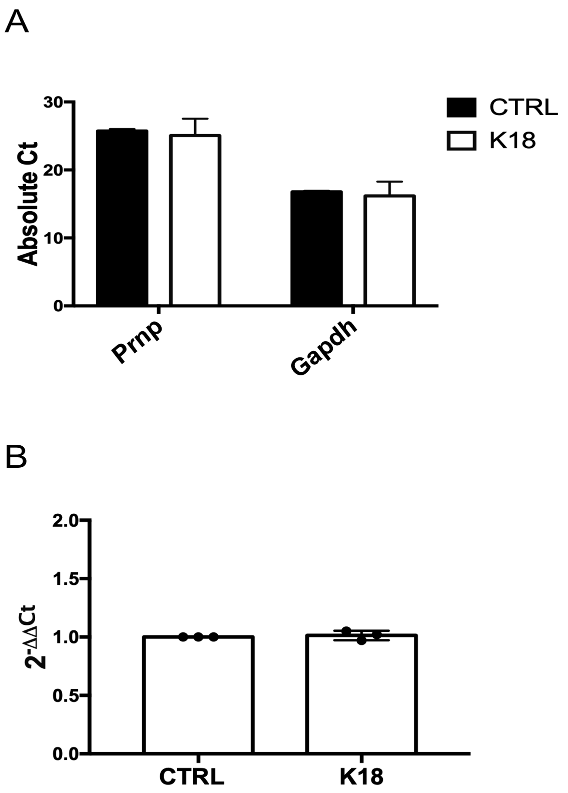


**Supplementary Figure S7. RT-PCR analysis of Prnp expression levels on control and treated N2a cells.** (A) Average values with SD of absolute CTs among the samples for *Prnp* and GAPDH genes. Every sample was analysed in duplicate. (B) Relative expression levels of *Prnp* gene normalized against GAPDH in control and K18-treated cells. The relative expression in control cells has been normalized to 1.

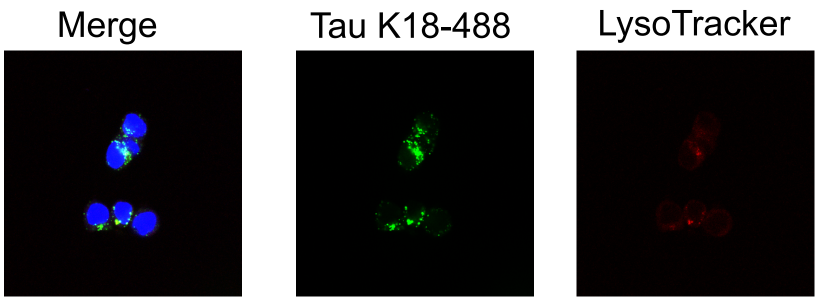


**Supplementary Figure S8. Immunofluorescence of ScN2a cells incubated with 2 uM of tau K18 amyloids (green) for 72 hours.** Nuclei are labelled in blue, while lysosomes were stained with LysoTracker (red). Images show the XY projection of one of the central sections of the stack, both as separate channels and as a merge of the three channels.

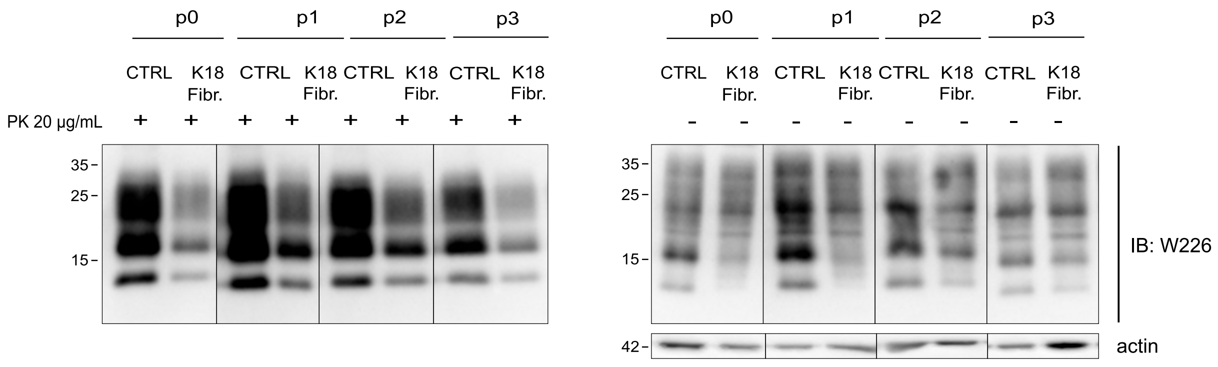


**Supplementary Figure S9. Western blot analysis of PrPSc in ScN2a RML cell lysates after treatment with tau K18 fibrils.** Following the treatment with tau K18 aggregates for 6 days, total PrP and PK-resistant PrPSc proteins were separated by SDS-PAGE and detected with anti-PrP Ab (W226). β-actin is a loading control.
